# Genetic evidence that the latency III stage of Epstein-Barr Virus infection is a therapeutic target for Multiple Sclerosis

**DOI:** 10.1101/390609

**Authors:** Ali Afrasiabi, Grant P. Parnell, Nicole Fewings, Stephen D. Schibeci, Monica A. Basuki, Ramya Chandramohan, David A. Brown, Sanjay Swaminathan, Fiona C. McKay, Graeme J. Stewart, David R. Booth

## Abstract

Genome wide association studies have identified >200 susceptibility loci accounting for much of the heritability of Multiple Sclerosis (MS). Epstein Barr virus (EBV), a memory B cell tropic virus, has been identified as necessary but not sufficient for development of MS, with evidence for disease causation. The molecular and immunological basis for this has not been established. LCL proliferation is driven by signalling through the EBV produced cell surface protein LMP1, a homologue of the MS risk gene CD40. We show that the CD40 ligand, CD40L, potentially through competitive signalling with LMP1, reduces LCL proliferation (p<0.001). The MS risk variants of the LMP1 signalling inhibitor, TRAF3, had altered expression in B cells and LCLs. Both CD40 and TRAF3 risk SNPs are in binding sites for the EBV transcription factor EBNA2. We have investigated transcriptomes of B cells and EBV infected B cells at Latency III (LCLs) and identified 47 MS risk genes with altered expression, associated with the risk genotype. Overall these MS risk SNPs were overrepresented in target loci of the EBV transcription factor EBNA2 (p<10^−16^), in genes dysregulated between B and LCLs (p<10^−5^), and as targets for EBV miRNAs (p<10^−4^). The risk gene ZC3HAV1 is the putative target for multiple EBV miRNAs. It amplifies the interferon response, and was shown to have reduced expression in LCLs for the risk allele. These data indicate targeting EBV EBNA2, miRNAs, and MS risk genes on the LMP1/LMP2 pathways, and the pathways themselves, may be of therapeutic benefit in MS.

## Introduction

Epstein Barr virus (EBV), a memory B cell tropic virus, has been identified as necessary but not sufficient for development of MS (*1*). This implies it’s necessary to initiate pathogenesis, but the molecular and immunological basis for this has not been established. Its potential importance in pathogenesis has been highlighted by the success of therapies removing memory B cells (antiCD20 monoclonal antibodies), and the failure of therapies which increase them (antiTACI monoclonal antibodies) (*2*). Causality is also indicated from three large studies using different approaches. From a large longitudinal study, adults without EBV antibodies who later developed MS also were infected by EBV (*3*). The risk of developing MS for those with the HLA-DRB1*1501 allele and for EBV infection is higher for the combination than due to each factor independently, suggesting an interaction and causality (*4, 5*). Finally, the time to developing MS is shorter for those with late EBV infection than those with early EBV infection, consistent with infection affecting development of disease (*6*).

First infection with EBV at a later age confers greater risk of developing MS than early infection (*7*). In addition, association studies indicate higher EBV loads for those with MS compared to controls (*8*), and increased T cell response to EBV antigen on therapy in those with a clinical response (*9*), suggesting an effect on disease progression. Due to its ability to immortalise B cells, EBV can cause a number of malignancies due to lymphomas (*10*), notably nasopharyngeal cancer and Hodgkin’s lymphoma. In immunocompromised individuals, such as those receiving transplants, unrestrained EBV infection can be fatal. Collectively these data indicate this near-ubiquitous virus is usually controlled by a sustained immune response, and are consistent with the hypothesis that failure of EBV control can induce disease, including MS. Moreover, EBV control may continue to be impaired and worsen in established MS. With increasing disease duration, EBV-specific T cells progressively decline, consistent with T-cell exhaustion and inversely proportional to anti-EBNA-1 titres (*11*). There is a higher incidence of spontaneous transforming events in long term culture of MS B cells (*12*), together supporting the existence of a higher proportion of latently EBV-infected B cells in MS.

Where EBV causes tumours, symptoms of infection, or uncontrolled lymphocyte proliferation, possible treatments include antiCD20 antibodies to remove the host target cell, anti EBV T cells from tissue banks, or genetic manipulation of anti EBV T cells (*13*). Vaccines to specifically boost immune response are in development (*13*). To promote further development of these approaches, and to identify novel ones, a better understanding of immune evasion by EBV is needed. The genetic variation that increases risk of MS may indicate molecular pathways controlling EBV, and so targets for improved therapy.

More than 200 gene loci have been associated with MS using genome wide association studies (GWAS) (*14*). For one of the first genes to be identified as an MS risk factor using GWAS, CD40 (*15*), a T cell activation gene, the protective allele was shown to have higher expression (*16*). This is counter-intuitive, since T cell activation is thought to promote MS risk. A potential explanation for this paradox is that higher expression of CD40 reduces signalling through the EBV protein LMP1, a homologue of CD40, inhibiting EBV survival in B cells. This could occur via competition for intracellular signalling molecules shared between these pathways. Increased cross-linking of CD40 has been shown to inhibit LCL proliferation (*17, 18*). Here we have investigated CD40L inhibition of LCL proliferation; and the effect of the CD40 risk genotype and that of another MS risk gene, TRAF3, a ligand for both CD40 and LMP1. We then sought other MS risk genes which might affect LMP1 and other Latency III signalling pathways in LCLs, using systematic and agnostic approaches.

EBV has four transcriptomes corresponding to three latency phases and a lytic phase. Latency III phase is the major target of the four phases for the immune response in chronic disease (*19*). To find risk genes potentially affecting control of EBV infection, firstly, we identified MS risk genes expressed differently in B cells and EBV infected B cells at Latency III (called LCLs, for lymphoblastoid cell lines) using RNAseq. Second, we used *in silico* data on the association of genotypes with expression in whole blood and LCLs. Third, we identified which of the most proximal genes to EBV transcription factor binding sites in LCLs were MS risk genes. Fourth, we used viral miRNA prediction software to identify putative EBV miRNA targets in MS risk genes dependent on MS risk genotype. We then tested the effect of risk genotype on expression of candidate genes in LCLs and B cells. These data provide genetic support for a facilitative role of EBV infection in MS, but do not prove it. They indicate molecular processes important in regulating LCL proliferation, and so molecular targets for control of EBV infection and potentially reducing MS progression.

## Results

### CD40 and TRAF3 expression and risk genotype in B cells and LCLs

Expression of CD40 is highest for the protective genotype (SNP rs1883832 T) in blood (*16*) and B cells (*20*). We hypothesized this higher expression decreased signaling through LMP1 through competition for the same signaling molecules. We first confirmed that its expression was higher in LCLs in GTEx data (*21*) for whole blood (n=396, p<10^−8^, rank of p value 29/4566 genes) and LCLs (n=117, p<0.06, rank 330/4566), and in the RTeQTL data (*22*) (n=955, combined p<10^−8^ for GTEx and RTeQTL data - see below). Earlier we had shown that the protective allele decreases sCD40 mRNA (exon 6 spliced out) in B cells and dendritic cells (*20*). In our LCL cohort (Westmead Institute for Medical Research; WIMR LCLs) the full length isoforms make up 80% of transcripts, whereas they are only 65% of transcripts in B cells (p<10^−9^, Figure 1A, B). Isoform usage was genotype independent in both cell types. We then tested if signaling through CD40 reduced LCL proliferation by culture with/without CD40L. We found that proliferation of LCLs in the presence of CD40L is decreased (p<0.0001, Fig 1C), with more inhibition for the protective genotype (p<0.03, Fig 1D). This is consistent with the CD40 protective genotype effect on MS being due to reduced susceptibility to Latency III proliferation of EBV.

**Figure 1.**
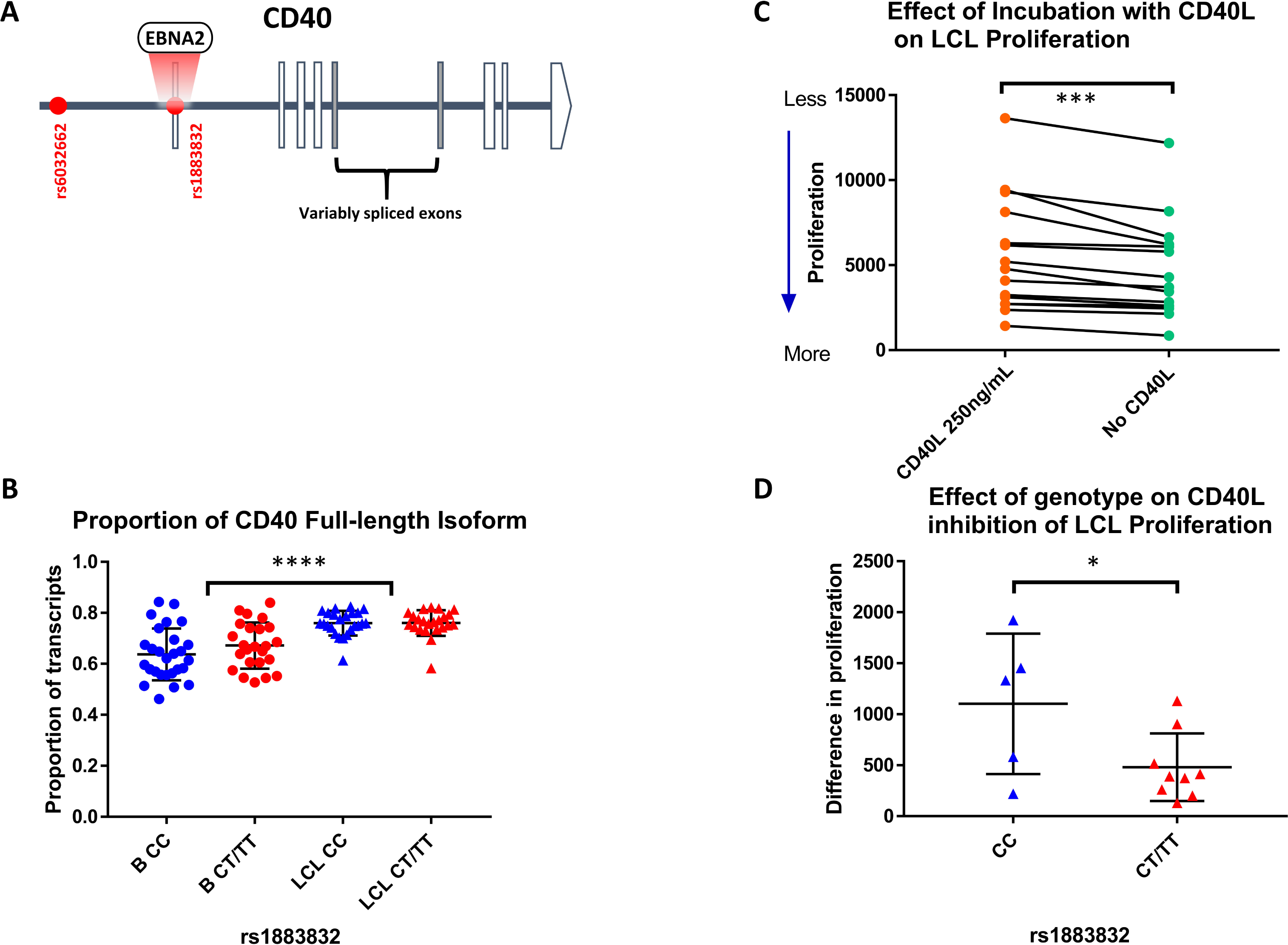
MS risk gene CD40 expression and signalling in B cells and LCLs. (A) Risk SNP rs1883832 is a binding site for the EBV transcription factor EBNA2. (B) The proportion of full length CD40 mRNA is higher in LCLs compared to CD19+ B cells, independent of rs1883832 genotype. (C) CD40L inhibits LCL proliferation (D) The inhibitory effect of CD40L on LCL proliferation is greater for the protective genotype (CC).

TRAF3 is an MS risk gene with two independent association signals, for SNPs rs12588969 and rs12147246. TRAF3 protein binds to the CTAR1 domain of LMP1, and reduces signaling through TRAF6, which binds to the CTAR2 domain of LMP1 (see Fig 7). We found that the protective variant of the risk SNP rs12588969 is associated with lower expression of TRAF3 in GTEx LCLs (p<0.0001) and in WIMR B cells, with a trend in LCLs (Fig 2), and replicated in the RTeQTL dataset (combined p<10^−15^), but not in blood. This SNP is in a locus bound by the EBV transcription factor EBNA2 (Fig 2A). The protective allele of the second SNP rs12147246 (using SNP rs12148050 as a proxy SNP) was associated with higher expression of TRAF3 in B cells and in blood, but not in GTEX or WIMR LCLs. The protective SNP of rs12148050 was associated with reduced proliferation of LCLs in the presence of CD40L (p=0.02), but no association was seen for rs12588969.

**Figure 2.**
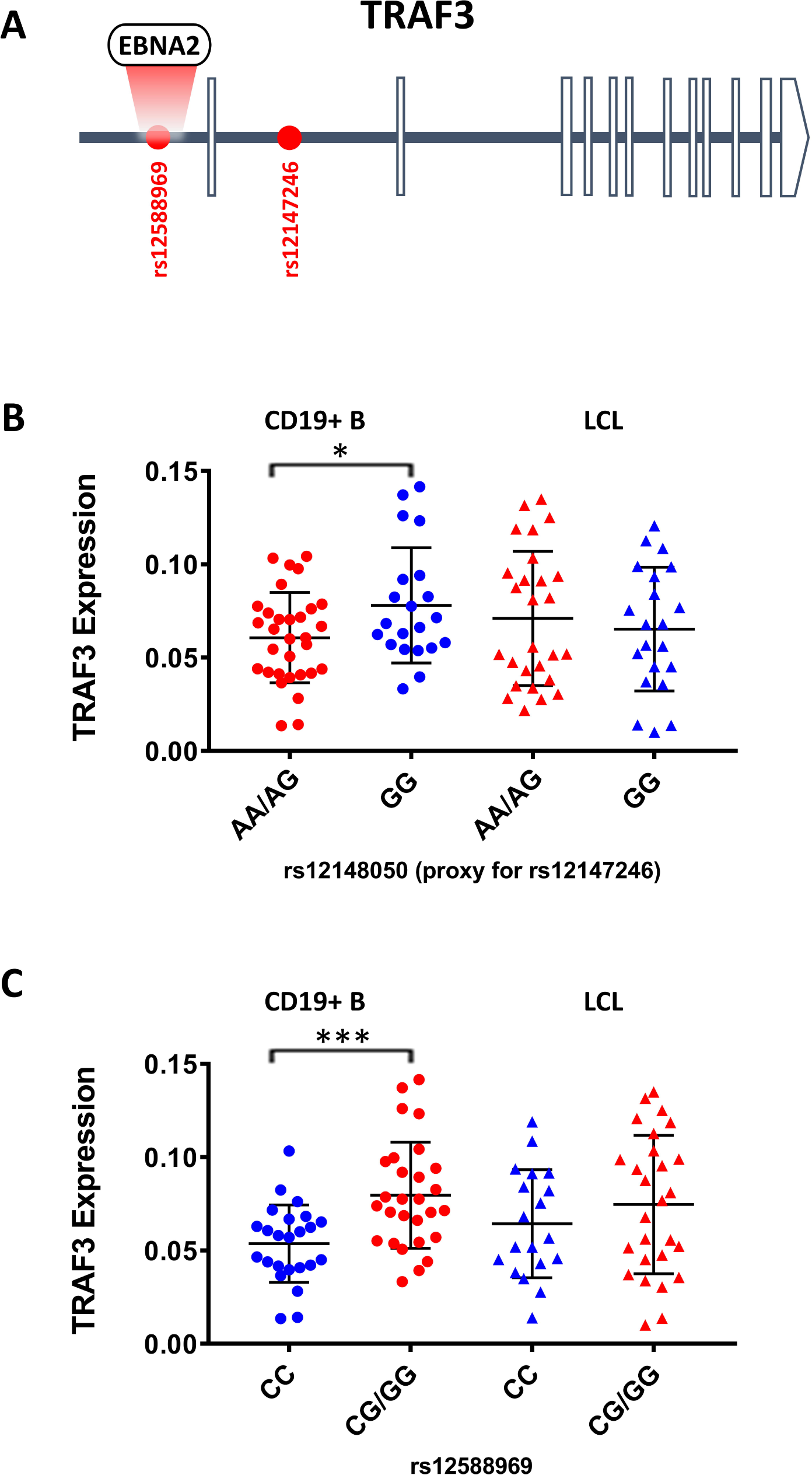
The effect of MS risk SNP genotype on the expression of TRAF3. (A) Risk SNP rs12588969 is a binding site for the EBV transcription factor EBNA2. TRAF3 expression is genotype dependent in CD19+B cells for MS risk SNPs rs12147246 (measured by proxy SNP rs1214050) (B) and rs12588969 (C).

These data are consistent with both CD40 and TRAF3 rs12148050 protective SNPs reducing susceptibility to EBV due to genotype effects in B cells and LCLs; conferring protection through decreasing signaling through the LMP1 pathways in newly infected B cells or LCLs. However the expression pattern for rs12588969 is inconsistent with this. The complexity of these findings for TRAF3 may be due to different isoform usage, use of the NFKB alternative pathway, or context dependent function in B and LCLs.

### The LCL Transcriptome

To identify other MS risk genes likely to contribute to variation in regulation of EBV infection we first screened for risk genes with altered regulation in EBV infected B cells (LCLs) compared to B cells. We used RNAseq to interrogate expression in *ex vivo* CD19+ B cells, and in LCLs derived from them using EBV strain B95.8 infection. Consistent with the different phenotypes, the transcriptomes were very different between infected and uninfected B cells. At a false discovery rate (FDR) of 0.01, 9017 genes were expressed differently (Fig 3A, 3B) (Supplementary Table 1). Differentially expressed genes were enriched for interferon stimulated genes (p<0.001, Fig 3C). Additional pathways over-represented included NF-kappaB signaling, apoptosis, cell cycle and viral processes (Fig 5B).

**Figure 3.**
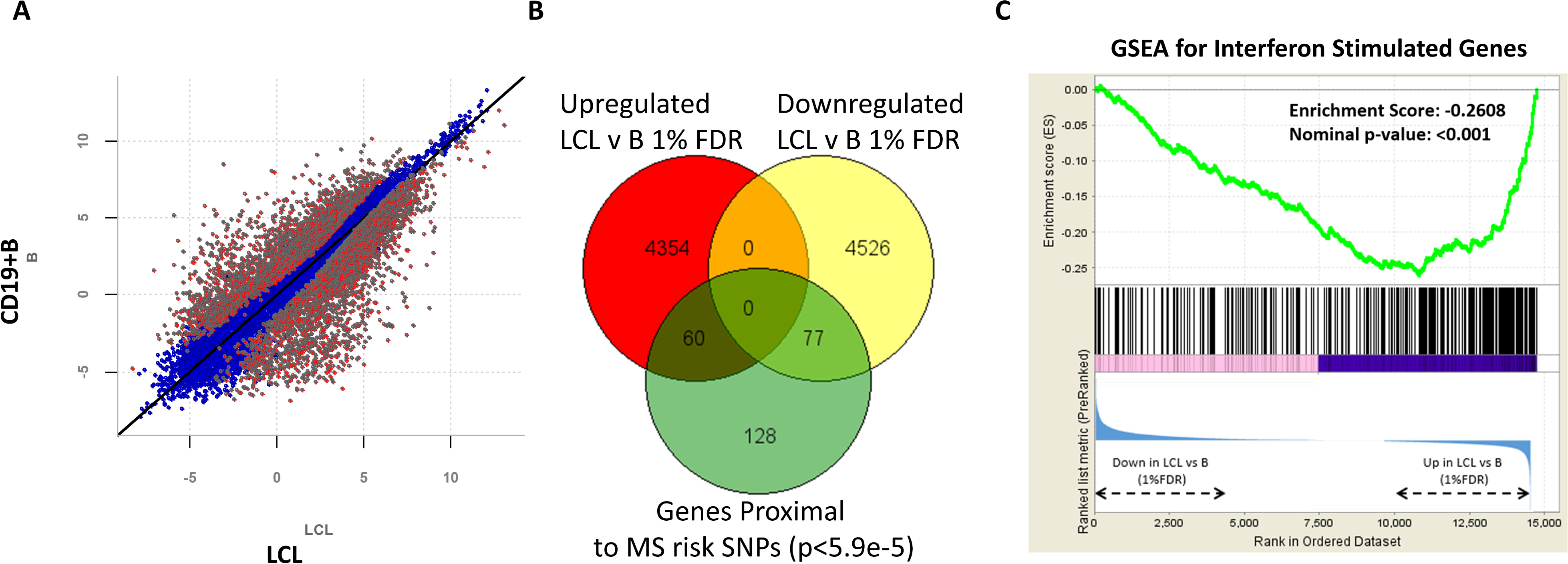
RNAseq reveals vast gene expression changes in LCLs compared to CD19+B cells. (A) Scatter plot highlighting genes differentially expressed at 1% FDR (red), (B) Genes proximal to MS risk SNPs are over-represented in the differentially expressed genes, (C) Gene Set Enrichment Analysis (GSEA) shows interferon stimulated genes are over-represented in genes upregulated in LCLs.

**Figure 4.**
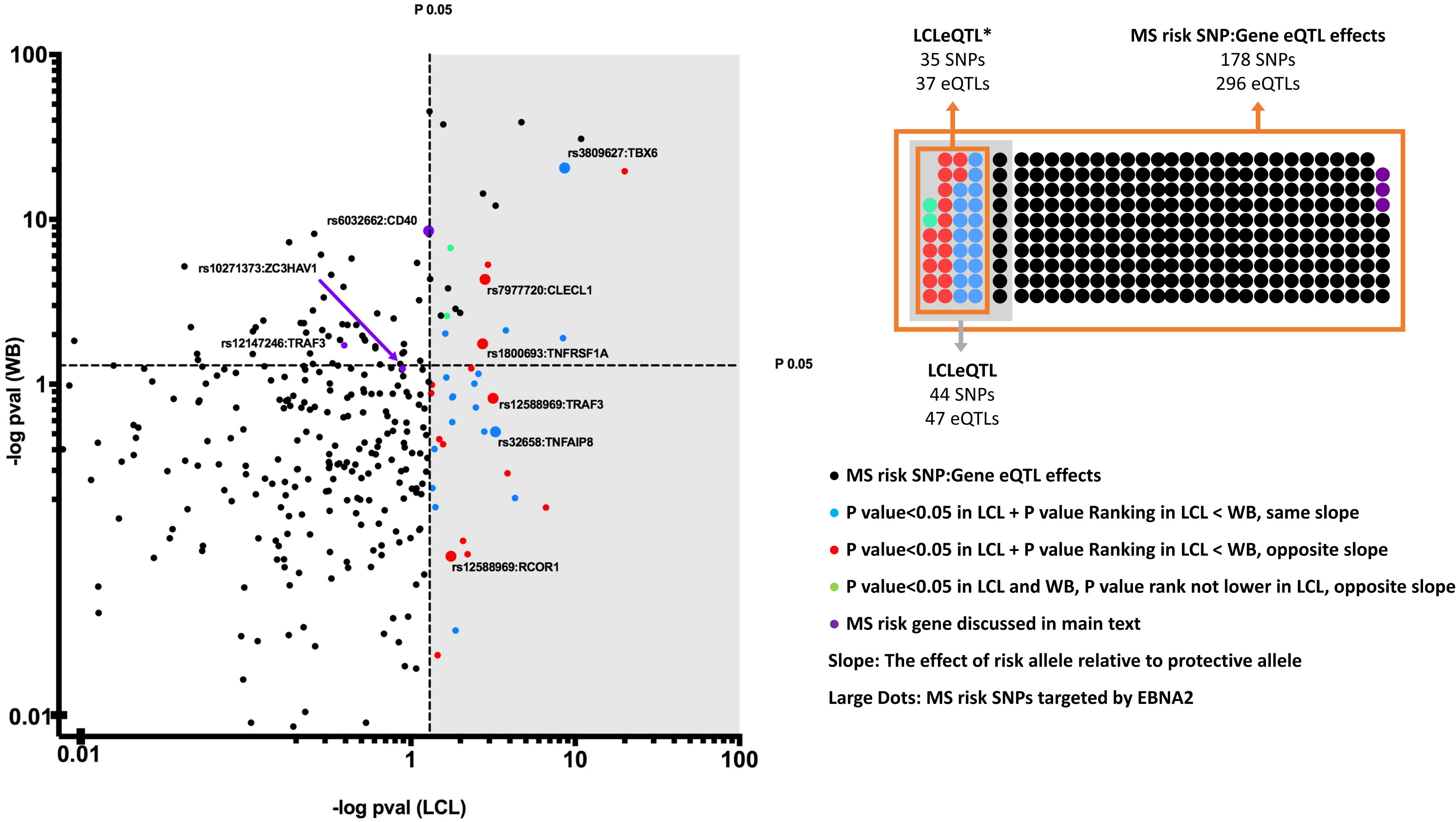
The effect of MS risk SNP genotype on expression of proximal genes in whole blood and LCLs. The GTEx eQTL dataset was first filtered for MS risk SNPs, and SNP:gene pairs were then plotted for effect of genotype (restricted to the genes closest to the MS risk SNPs – the proximal genes). SNP:gene pairs that were more strongly associated with expression in LCLs (coloured blue) and those with a different risk allele effect in LCLs compared to whole blood, more significant in LCLs (red) or not (green) were identified as LCLeQTL*.

**Figure 5.**
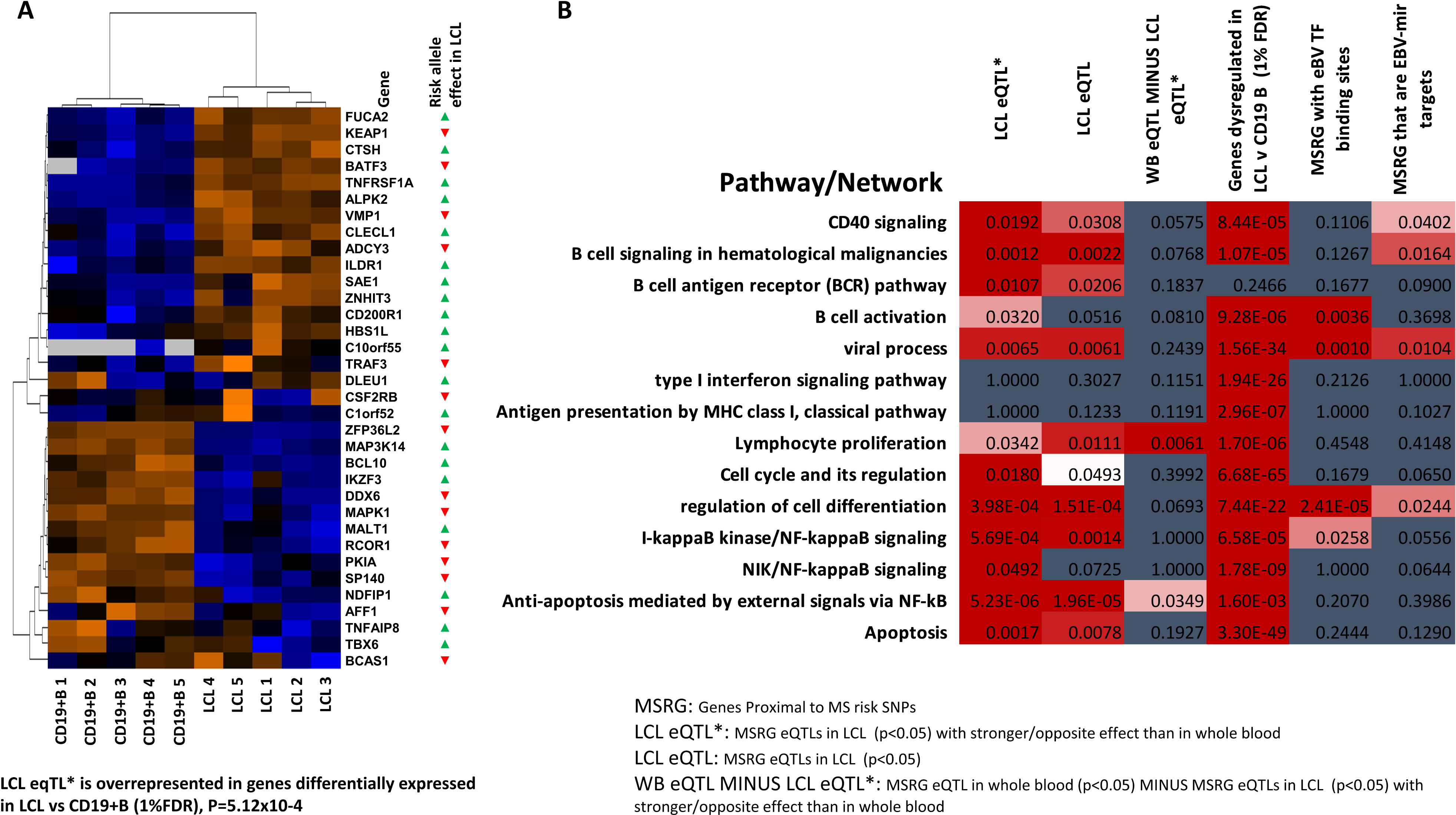
Genes and pathways with a stronger association of expression in LCLs compared to whole blood. (A) Heat map showing expression level of genes corresponding to the SNP:gene pairs that were more strongly associated with expression in LCLs compared to whole blood (LCLeQTL*) in CD19^+^ B cells and LCLs. (B) Over-representation of pathways and networks in; LCLeQTL*(MSRG eQTLs in LCL (p<0.05) with stronger/opposite effect than in whole blood); LCLeQTL(MSRG eQTLs in LCL (p<0.05)); WB eQTL minus LCLeQTL*(MSRG eQTL in whole blood (p<0.05) minus MSRG eQTLs in LCL (p<0.05) with stronger/opposite effect than in whole blood); genes dysregulated in LCLs vs CD19^+^ B cells at 1% FDR; MSRG with EBV transcription factor binding sites; MSRA that are EBV encoded miRNA targets. MSRG, Genes proximal to MS risk SNPs (*14*). TF, transcription factor.

**Figure 6.**
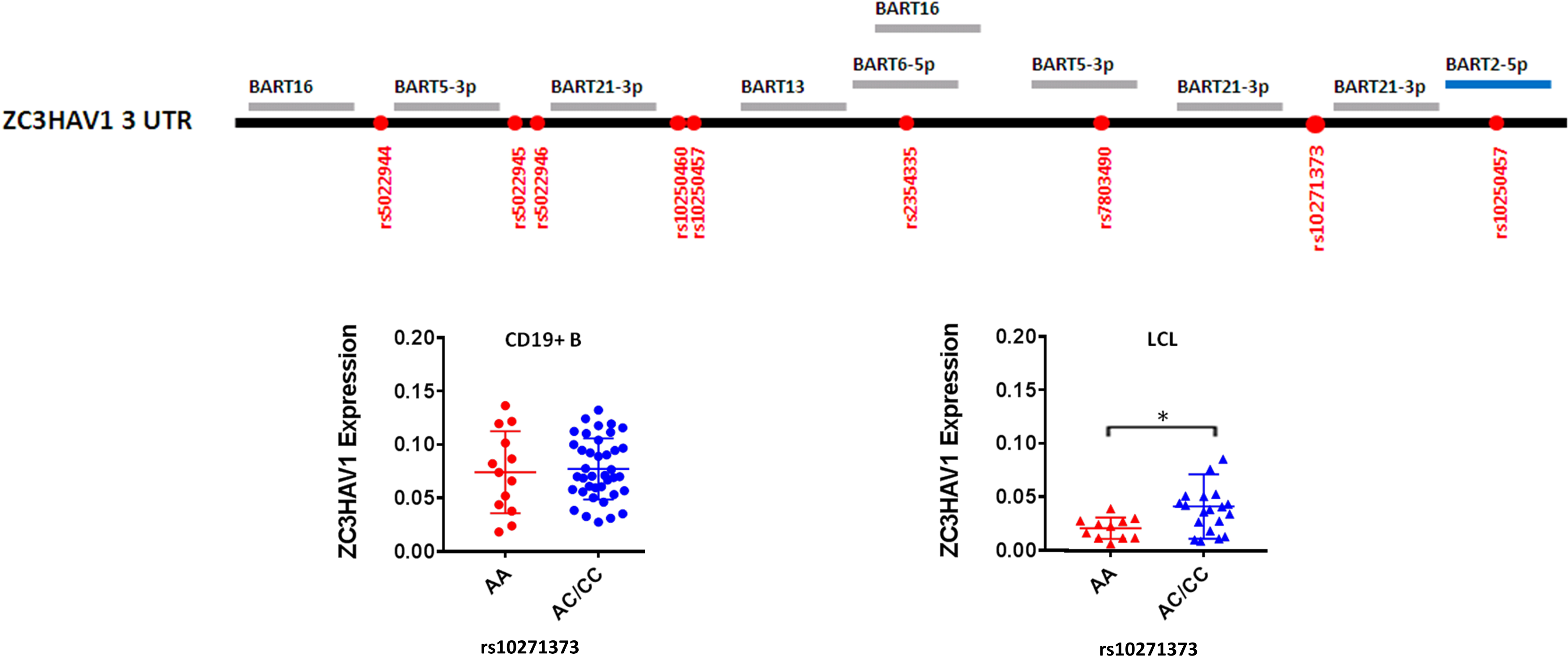
MS risk gene ZC3HA is a putative target for EBV encoded miRNA and expression is genotype-dependent in LCLs but not B cells.

**Figure 7.**
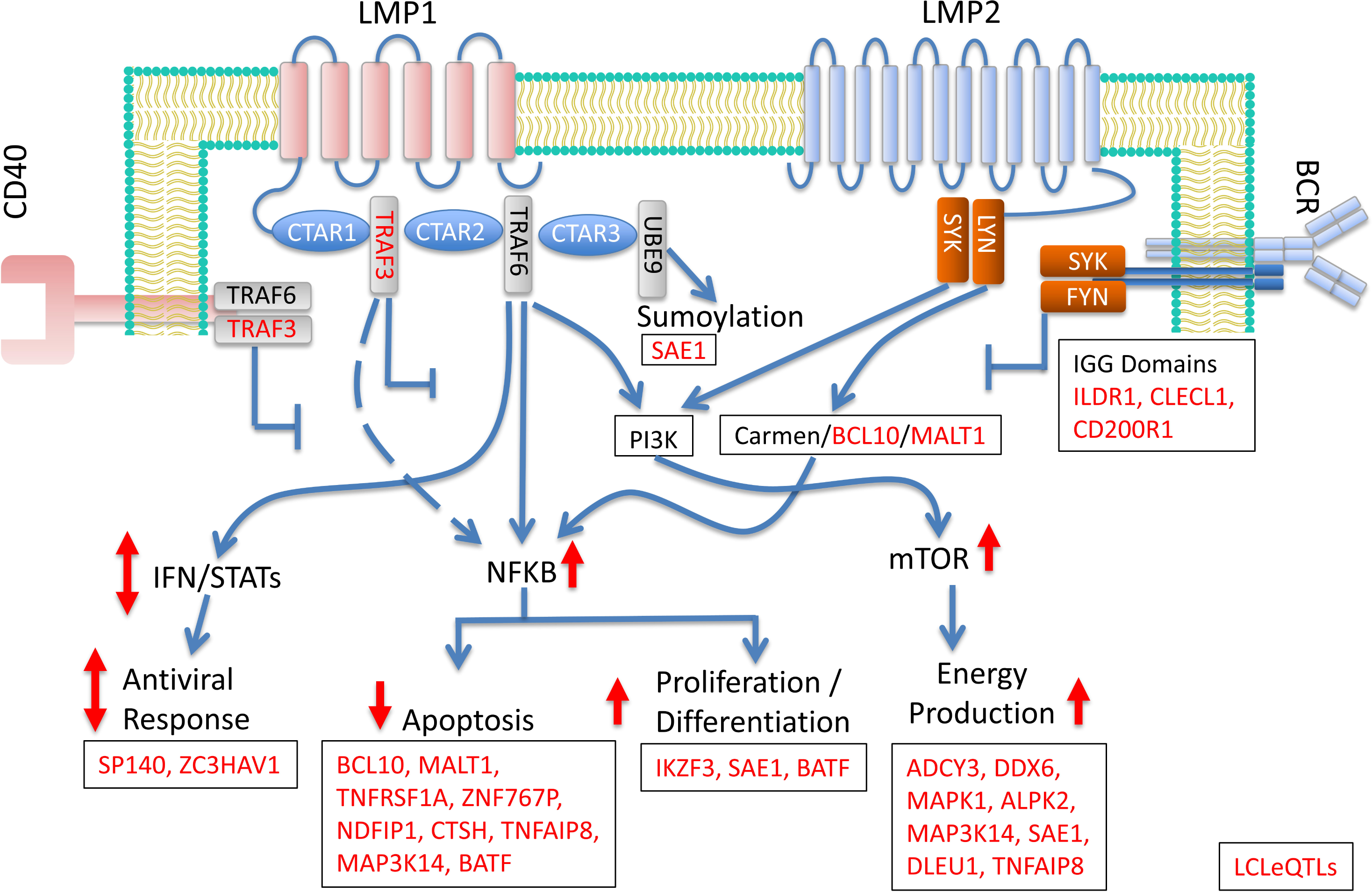
Model of LCLeQTL* gene roles in Latency III signaling pathways. Signaling from EBV proteins LMP1 and LMP2 leads to upregulation of NFKB, sumoylation, mTOR1 pathways; and altered IFN regulation. LCLeQTL* genes with roles on these pathways are in red. Signaling from the LMP1 homologue and MS risk gene CD40 inhibits LMP1 function (see Fig 1). For similar reasons, signaling through the BCR, in this model, blocks LMP2 pathways.

### MS risk genes that are LCL eQTLs

Since risk SNPs mainly affect pathogenesis by altering gene expression (IMSGC, 2015), we identified SNPs associated with gene expression (expression quantitative trait loci; eQTLs) in LCLs (LCLeQTLs, n=47 of 265, p<0.05, Supplementary Table 2) using GTEx data as a discovery dataset, and the RTeQTL data (*22*) as a replication set. Of the 47 GTEx LCLeQTL SNPs, 31 were tested in at least one of the RTeQTL cohorts, and 28 of these were replicated at p<0.05, mostly with much lower p values (Supplementary Table 3). We then identified those 47 GTEx LCLeQTL SNPs more associated with expression in LCLs than whole blood, as this would favour LCL expression as driving the pathogenic basis for their association with MS, more than expression in the immune cells of the blood (Fig 4). As the statistical power for the whole blood cohort was greater, we based this comparison instead on rank of p value, rather than raw p values. Of these, 18 had the same genotype effect on expression in LCLs and blood, 17 of these had opposite genotype associations with expression. Finally, two did not have lower p values in LCLs, but are included in the list as having opposite genotype effects between LCLs and blood. This list of 37 SNP:gene pairs we call the LCLeQTL* (Supplementary Table 4). 33 of 47 LCLeQTLs were in the genes differentially expressed between B cells and LCLs (over-representation p=3.83×10^−5^), and 24/37 of the LCLeQTL* genes were (p=5.12×10^−4^). Of the blood eQTL MS risk genes (Supplementary Table 5), a lower proportion were in the differentially expressed gene list (33/57, p<0.001), even less with the LCLeQTL genes removed from the blood eQTLlist (Supplementary Table 6, p=0.007). Of the 265 MS risk genes, 139 were differentially expressed between LCL and B cells (p=5.99×10^−5^). Despite these over-representations of LCLeQTL SNPs, blood eQTLs and especially MS risk genes in the genes differentially expressed between EBV infected and B cells, it remains possible that the enrichment may point to the B cell functions of these genes/SNPs rather than viral contribution to pathogenesis.

### MS risk genes potentially regulated by EBV transcription factors or miRNAs

To identify EBV-specific enrichment, we sought risk SNPs/genes co-located with EBV transcription factor binding peaks. There were 871 EBV transcription factor (TF) peaks identified in the genome (*23*). MS risk SNPs were co-located (28 of 201 mapped) with EBV TF binding peaks at an extraordinary level of enrichment (Supplementary Table 7, p= 3.17×10^−16^). Of the 47 LCLeQTLs, 5 SNPs were co-located with EBNA2 peaks (p=1.83×10^− 4^). The proximal genes to these 5 SNPs were TRAF3, RCOR1, TBX6, TNFAIP8, TNFRSF1A, and CLECL1. These 5 SNPs were also in the 37 LCLeQTL*s list (p=5.1×10^−4^). We then sought genes which might be regulated by EBV miRNAs using the pipeline described in Supplementary Figure 5 using the application RepTar, which is designed to predict mRNA binding by viral miRNAs, for the miRNA expressed in latency III by strain B95-8. 29 binding sites were identified within 30 base pairs of MS risk SNPs (or SNPs in LD > r^2^ 0.8, Supplementary Table 8). One gene, ZC3HAV1, with a p value of 0.12 in the GTEx data, had 13 putative EBV miRNA target sites in its 3’end (Supplementary Table 10). One of these recognition target sites, for EBV miRNA BART2-5p, covers one SNP (rs10250457) in LD with the risk SNP rs10271373. The risk gene ZC3HAV1 is known to be used by the host to amplify the interferon response, so the higher expression for the protective allele is consistent with increased IFN response and so decreased immune evasion by EBV.

Further, from a study using experimental validation of miRNA/target duplexes, the PAR-Clip data (*24*), of 265 risk genes, 31 had putative EBV miRNA targets (over-representation p=6.7×10^−4^) (Supplementary Table 9); of 47 LCLeQTLs 11 were targets (p=3.82×10^−5^); of 37 LCLeQTL* 10 were targets (p=0.015); and of the 57 blood eQTLs minus these LCLeQTL* 6 were targets (ns).

### MS Risk Genes on the LMP1/LMP2 Signalling Pathways

Finally, the LCLeQTL*s were mapped on to the LMP1 and LMP2 signaling pathways (Fig 7). The LMP1 pathway was as defined previously (*25*), which overlaps the CD40 pathway as described (*26*). This pathway was confirmed and extended in the work of Li et al (*27*), who used nasopharyngeal tumour mutations, a semi-agnostic approach, to define the pathway. The LMP2 pathway was as defined in Cen and Longnecker (*28*), who described its overlap with the B cell receptor (BCR) signaling pathway. LCLeQTL* gene-encoded protein TRAF3 binds directly to LMP1 and CD40; IgG domain containing proteins (genes CLECL1, CD200R and IL1DR) may bind or compete with LMP2/BCR. Molecules one signaling node down from this affect sumoylation (SAE1) and NFKB activation via the BCR (MALT1 and BCL10). NFKB is activated by both LMP1 and LMP2 pathways, and in turn regulates genes controlling apoptosis (different roles for BCL10 and MALT1 than above; and TNFRSF1, ZNF767P, NDFIPP1, CTSH, TNFAIP8, MAP3K14), proliferation and differentiation (IKZF1, SAE1). The LCLeQTL* transcription factor BATF interacts directly with NFKB in LCLs (*29*), and TNFRSF1A directly activates it (*30*). The mTOR complex is activated by both LMP1 and LMP2, and enables the vast changes in energy production required for the proliferating B cells in latency III (*31*). The LCLeQTL* genes regulating ATP are ADCY3, DDX6, MAPK1, ALPK2, MAP3K14, SAE1, DLEU1 and TNFAIP8 (GO annotation). Finally, the interferon pathway is dysregulated in latency III by altering expression of transcription factors such as IRF5, IRF7 and the STATs (*32*). This may underpin the genetic associations with expression seen for ZC3HAV1 and SP140 in LCLs, so that EBV contributes to altered expression to dampen the IFN response. Several LCLeQTL*s were transcription factors of undefined roles: ZNHIT3, TBX6, RCOR1, ZFP36L2, AFF1. These are likely to mediate some of the global changes to B cell function on infection. Also, several LCLeQTL*s had functions not readily attributable to LMP1/LMP2 signaling; and for others their functions are largely unknown. Collectively, there is strong support for LCLeQTL* genes affecting latency III gene expression programs.

## Discussion

From the recently expanded list of genes affecting MS susceptibility (*14*), we sought evidence that response to EBV latency III infection contributed to susceptibility to MS. Expression of risk genes, the LMP1 homologue CD40 and the LMP1 ligand TRAF3, was affected by risk genotype in EBV infected cells and B cells. Infected B cell proliferation was reduced on signaling through CD40, and more so for the protective genotype. The number of MS risk SNPs associated with genes differentially expressed between infected and uninfected B cells was 137 of 265, much more than would be expected by chance. The number of MS risk SNPs where the genotype was associated with proximal gene expression (47 of the 265 risk SNPs) in LCLs was also much higher than would be expected by chance. 37 of these 47 were more associated with expression in LCLs than they were with blood. EBV transcription factor binding sites are overrepresented among MS risk genes and among LCLeQTLs, including at risk genotype sites for 6 genes. EBV miRNA predicted binding sites are over-represented among MS risk SNPs/genes, genes dysregulated on EBV infection, and LCLeQTLs. EBV miRNA BART2-5p is predicted to bind the interferon response gene ZC3HAV1 at the risk SNP, and the protective variant of the risk SNP for this gene had reduced expression in LCLs. Finally, the 37 MS risk genes identified as LCLeQTLs have plausible roles in signaling on the LMP1 and LMP2 pathways in EBV latency III.

Although these data provide genetic evidence that EBV has a facilitative role in MS, they are not conclusive. The association of MS risk genotypes with LCL expression, and the enrichment of risk genes involved in B cell proliferation and utilized by EBV, may be due to B cell processes contributing to disease, and independent of the role in EBV latency III. Also, even though the interaction between CD40/TRAF3/LMP1 would predict protective genotypes decrease EBV latency III proliferation, this proliferation may be independent of the pathogenic effect in MS of these genes. Finally, the association of MS risk SNPs with expression and LCL proliferation may be different in B cells from people with MS.

Genes affecting MS susceptibility favour processes leading to dysregulated immune responses, and if the processes are ongoing and drive MS pathogenesis, reversing these would be expected halt progression. Monoclonal antibodies to risk gene IL2Ra (drug Daclizumab) and to CD49D, ligand of risk gene VLA4 (drug Natalizumab) are effective therapies for MS (*2*). More generally, drugs which remove immune cells expressing risk genes, such as antiCD20 (B cells) and antiCD52 (Alemtuzumab); or which corral them in secondary lymphoid organs (Fingolimod) are also effective.

Similarly, if poor regulation of EBV infection contributes to MS susceptibility and progression via processes tagged by the MS risk gene LCLeQTLs, therapeutic strategies to favour immune control of EBV may halt or slow progression. Especially attractive are targets on the EBV genome, since these greatly increase specificity for EBV compared to the host genes which regulate LMP1 and LMP2 signaling. Of these EBV targets, methods of reducing EBNA2 expression or activation through its binding partners are in development for other conditions. Farrell et al (*33*) have shown a cell-permeable peptide inhibiting EBNA2 binding to co-transcription factor CBF1 results in downregulation of EBV proteins LMP1 and LMP2 and reduced LCL proliferation. EBV miRNAs such as miRNA BART2-5p, implicated here as regulating the interferon response through ZC3HAV1, could be inactivated with complementary nucleic acid therapeutics, as currently used for familial amyloidotic polyneuropathy (*34*).

Although it is challenging to prove the MS risk genotype is associated with MS because it alters Latency III proliferation and immune evasion, further support would come from demonstrating (1) that reduction of EBNA2 and targeted EBV (B95.8 strain) miRNAs reduces the genotype association with expression in LCLs; (2) altered expression of the risk gene corresponding to the LCLeQTL effect alters LCL proliferation, especially through the pathway the gene affects; and (3) effects may be exaggerated in LCLs derived from MS patients. Testing of genotype effects on expression and function in other EBV phases is also warranted. Finally, the killing of LCLs by EBV specific T cells or NK cells in co-cultures may be risk genotype and gene dependent.

These data indicate many genetic risk factors identified in genome wide association studies for MS susceptibility have roles consistent with a dysregulated response to EBV infection, and so in this way contribute to MS pathogenesis. They point to particular molecular processes important in regulating LCL proliferation, and so molecular targets for control of EBV infection to potentially reduce MS progression. Specifically, these data indicate targeting EBV EBNA2, EBV miRNAs, and MS risk genes functioning on the LMP1/2 pathways, and the pathways themselves, may be of therapeutic benefit in MS.

## Methods

### Samples

All blood samples were collected from healthy controls with informed consent (Westmead Hospital Human Research Ethics Committee Approval 1425). B lymphocytes were purified by immunomagnetic human B cell enrichment Kit (Stem Cell Technologies) according to manufacturer's instructions. For LCL generation, fresh or frozen PBMCs were infected with supernatant from B95.8 producer cell line for 1 hour at 37°C. Cells were then suspended in complete medium, consisting of RPMI-1640 (Lonza) supplemented with 10% fetal bovine serum (FBS, Sigma Aldrich), 2 mM L-glutamine (Life Technologies), 50 units per ml penicillin/50 g per ml streptomycin (Life Technologies) plus 2 μg/ml of cyclosporin A (Sigma Aldrich) and plated at 2.5×10^6^ or 5×10^6^ cells/well in 48-well plates. Cells were fed weekly, expanded into 25 cm^2^ flask. LCLs were cryopreserved in 10% DMSO (MP Biomedical) 50% FBS and RPMI.

### CD40 proliferation

For proliferation readout, LCLs were first labelled with Cell Trace Violet (CTV, Life Technologies) at a final concentration of 5mM. 5×10^4^ LCLs were cultured with or without CD40L (250ng/mL, Adipogen) in 5mL polystyrene tubes (Becton Dickinson, BD) for 5 days. Day 5 cells were harvested and ran on FACSCantoII flow cytometer (BD). Median Fluorescence Intensity (MFI) of CTV was analysed using FlowJo.

### RNAseq

Global gene expression profiling using RNAseq was carried out for n=5 LCL and n = 5 CD19+ B cells. Total RNA was first isolated using the RNeasy Mini Kit (Qiagen) before RNAseq library preparation using the TruSeqV2 Library Preparation kit (Illumina). The indexed libraries were pooled 10 plex and 50bp single end reads were sequenced on the HiSeq 2500 (Illumina). Reads were assessed for quality using FastQC, aligned to hg19 using TopHat2 (*35*), and summarised to RPKM gene level expression using SAMmate (*36*). Differentially expressed genes were calculated using EdgeR (*37*). A cut-off of 1% false discovery rate was selected to select differentially expressed genes, see Supplementary Table 1.

### Genotyping and Gene expression

In the Westmead Cohort. CD40 (SNP rs1883832), TRAF3 (SNPs rs12588969, rs12148050), ZC3HAV1 (SNP rs1021373) were genotyped using Taqman Assays, and gene expression was assayed using Taqman probes probes Hs00912657_m1, Hs00936778_m1 and Hs01002915_g1 respectively (Life Technologies). Splicing of CD40 was determined as previously described (*20*).

### GTEx eQTL in LCL and WB

The rAggr website was used to prepare the variant call format (VCR) for all 201 MS risk SNPs from rs ID and genomic coordinates (199 SNPs were located). eQTL data for 187 of these 199 MS risk SNPs were extracted from the GTEX dataset, for LCL and whole blood (WB). The effect of risk allele relative to protective allele based on the slope data was then extracted. We filtered proximal genes from all eQTL data associated with MS risk SNPs in LCL and WB (Supplementary Figure 2) and merged the two datasets as a final SNP:Gene table to compare the genotype effect of each MS risk SNPs in LCL and WB. We identified the SNP:Gene eQTL pairs with higher association in LCL than WB based on the P value being less than 0.05 in LCL, and of a lower rank in P value ranking system for SNP:Gene list in LCL than WB. Also, of the genes with a p<0.05 in LCLs, those with a reverse slope between LCLs and WB were included. The set of MS risk SNPs VCF format, all eQTL data for MS risk SNPs in LCL and WB, eQTL data for proximal genes of MS risk SNPs and SNP:Gene pairs eQTL with higher association in LCL than WB and EBV are reported in Supplementary Tables 2-5, and the detailed workflow is Supplementary Fig 3.

### Pathway analysis

MetaCore was used to undertake pathway analysis for proximal genes which are resulted from: (I) EBV transcription factors binding to MS risk SNPs, (II) SNP:Gene eQTL pairs with higher association in LCL rather than WB, (III) EBV miRNA targets, (IV) all proximal SNP:Gene eQTL pairs in LCL, all proximal SNP:Gene eQTL pairs in WB and (V) all proximal SNP:Gene eQTL pairs in WB without SNP:Gene eQTL pairs with higher association in LCL rather than WB.

### EBV Transcription factors/MS risk SNPs

EBV transcription factors’ binding peak ChIP-seq data were extracted using the Regulatory Element Local Intersection (RELI) tool. Genomic coordinates of ChIP-seq data for BZLF1, EBNA2, EBNA3c, EBNA1 and EBNALP EBV transcription factors in LCL were prepared as a BED file. Bedtools v2.26.0 software was used to identify the overlap between MS risk SNPs, SNPs in LD with them, and EBV transcription factor binding peaks (Supplementary Table 7). All steps of this workflow are illustrated in Supplementary Fig 4.

### EBV miRNA/MS risk SNPs

The 44 EBV encoded miRNAs’ mirRBase name and 223 MS risk genes’ RefSeq accession number were extracted from miRBase (http://www.mirbase.org/) release 21.0 and Table Browser tool of UCSC genome browser (https://genome.ucsc.edu/cgi-bin/hgTables), respectively. The SNP list includes >200 original MS risk SNPs which is reported by IMSGC GWAS study, and 4693 SNPS in linkage disequilibrium (LD) with MS risk SNPs with MAF≥0.1 and R^2^≥0.8 threshold in the CEU population. The rAggr (http://raggr.usc.edu/) were used as a LD calculator tool using the Hg19 genome assembly. Putative EBV miRNA seed sites were predicted by RepTar (http://bioinformatics.ekmd.huji.ac.il/reptar/), using settings for minimal free energy of binding (MFE) ≤ −10, normalized minimal free energy of binding ≥ 0.1, G-U base pairs fraction ≤ 0.25, site conservation ≥ 0 and repeating motifs ≥ 1. Based on the RISC complex foot print on mRNA, we extended the start and end positions of genomic coordinates of predicted seed sites 30 bp as a flanking region. The set of overlaps between MS risk gene LD blocks and seed sites is reported in Supplementary Table 8. All steps of this workflow are illustrated in Supplementary Figure 5.

## Statistics

Analysis of gene expression differences in the WIMR LCL and B cell cohort was conducted using GraphPad Prism 7 (GraphPad Software, USA) using paired and unpaired (where appropriate) two-tailed T-tests to compare between groups. P values for overlaps were calculated using the following online tool: http://nemates.org/MA/progs/overlap_stats.

## Supplementary Tables

**Supplementary Table 1.** Genes dysregulated due to EBV infection

**Supplementary Table 2.** LCLeQTL gene list.

**Supplementary Table 3.** Replication and combined P value for LCLeQTL gene list

**Supplementary Table 4.** LCLeQTL* gene list.

**Supplementary Table 5.** Whole Blood eQTLs gene list.

**Supplementary Table 6.** Whole Blood eQTLs minus LCLeQTL*.

**Supplementary Table 7.** EBV Transcription Factor interactome gene list.

**Supplementary Table 8.** Predicted EBV miRNA targetome among MS risk genes (RepTar).

**Supplementary Table 9.** EBV latent miRNA targetome gene list (PAR-Clip data).

**Supplementary Table 10.** EBV miRNAs’ binding sites on ZC3HAV1 3UTR.

## Supplementary Figures

**Supplementary Figure 1.** CD40 isoform uasage in B cells and LCLs

**Supplementary Figure 2.** Volcano plot of association of MS risk SNPs with proximal gene expression in whole blood and LCLs.

**Supplementary Figure 3.** Pipeline to identify LCL eQTLs.

**Supplementary Figure 4.** Pipeline to identify overlap of EBV transcription factor binding peaks and MS risk loci.

**Supplementary Figure 5.** Pipeline to identify EBV miRNA targets in MS risk loci

## Acknowledgements

DRB was funded by NHMRC Senior Research Fellowship 1078494; GP by MSRA/JDRF/Macquarie Group Foundation Postdoctoral Research Fellowships, and AA by a Du Pré Grant from MSIF. Further support was received from NHMRC project grant 1065157; and MSRA incubator grants to SS, FM and DRB.

## Author Contributions

DRB, GP, AA, and FM conceived the project; GP and SDS conducted the transcriptomic experiment; AA, GP and DRB conducted the in silico analyses; SDS, NF, MB and RR conducted laboratory experiments with the WIMR samples; all authors contributed to discussions; funding supplied by grants to DRB, GS, DPB, SSw, and FM. DRB drafted the manuscript, reviewed by all authors.

## Abbreviations

EBV: Epstein-Barr virus
MS: Multiple Sclerosis
eQTL: expression quantitative trait loci
LCL: lymphoblastoid cell lines
GWAS: genome wide association study
TF: transcription factor

